# Mergeomics: integration of diverse genomics resources to identify pathogenic perturbations to biological systems

**DOI:** 10.1101/036012

**Authors:** Le Shu, Yuqi Zhao, Zeyneb Kurt, Sean Geoffrey Byars, Taru Tukiainen, Johannes Kettunen, Samuli Ripatti, Bin Zhang, Michael Inouye, Ville-Petteri Mäkinen, Xia Yang

## Abstract

Mergeomics is a computational pipeline (http://mergeomics.research.idre.ucla.edu/Download/Package/) that integrates multidimensional omics-disease associations, functional genomics, canonical pathways and gene-gene interaction networks to generate mechanistic hypotheses. It first identifies biological pathways and tissue-specific gene subnetworks that are perturbed by disease-associated molecular entities. The disease-associated subnetworks are then projected onto tissue-specific gene-gene interaction networks to identify local hubs as potential key drivers of pathological perturbations. The pipeline is modular and can be applied across species and platform boundaries, and uniquely conducts pathway/network level meta-analysis of multiple genomic studies of various data types. Application of Mergeomics to cholesterol datasets revealed novel regulators of cholesterol metabolism.

---

Most non-communicable diseases stem from a complex interplay between multiple genes and cumulative exposure to environmental risk factors ^1^. An emerging hypothesis regarding the underlying pathogenic processes is that exposure to genetic or environmental risk factors results in progressive and chronic regulatory perturbations to molecular and cellular processes that would otherwise maintain normal homeostasis. In recent years, the advance of omics technologies has greatly enhanced our ability to test this hypothesis with genome-scale molecular datasets that are also publicly available to the scientific community. Large-scale data repositories such as dbGaP for population-based genetic datasets ^2^ and Gene Expression Omnibus and ArrayExpress for gene expression and epigenomics datasets ^3, 4^ are continuously expanded with new experiments, data acquisition projects such as ENCODE and GTEx are generating multidimensional coherent datasets and the necessary basic framework to bridge the gaps between diverse genomics datasets ^5,6, 7^

An isolated omics study can provide only a partial view of the biological system. For example, a genome-wide association study (GWAS) can reveal the statistical associations between genetic loci and disease status, and implies causal effects, but pinpointing the causal genes, their corresponding causal variants and mechanisms has proven challenging ^8,9^. Additionally, evolutionary constraints restrict the ability of GWAS to detect central regulatory genes ^10^, and directly translating genetic associations of common variants with subtle effects may miss novel therapeutic targets. On the other hand, gene expression or epigenomic profiling can detect associations between disease and genes or epigenomic markers, but these associations are correlative in nature. By integrating different types of data, it becomes possible to circumvent the limitations of individual studies and better identify disease-causing DNA variants and their downstream molecular targets. For instance, when DNA and RNA are measured simultaneously, it is possible to determine if a particular genetic variant affects the downstream expression of a gene in a genetics of gene expression or eQTL analysis ^11,12^. Furthermore, if a genetic variant resides in a functional site associated with transcription factor binding, epigenetic modification, or protein regulation, as revealed by the ENCODE project ^13, 14^, it becomes possible to narrow in on the potential targets.

In parallel to large-scale genomic projects, new computational tools are required to convert massive genomics data into biological insights that can lead to novel mechanistic hypotheses. Pathway-based analysis tools such as MAGENTA ^15^ and iGSEA4GWAS ^16^ that integrate GWAS with curated pathways and methods that additionally take into consideration of eQTL information ^17^ have been developed. Beyond knowledge-based pathway analysis, data-driven approaches utilizing gene regulation and protein-protein interaction networks have been developed to identify the most likely pathological perturbations and target genes for disease-associated loci. Network biology aims to identify high-level regulatory patterns that characterize systemic functions with nodes representing genes or other molecular entities and edges representing the relations between nodes^18,19,20^. Networks can be classified into curated knowledge (e.g. metabolic pathways), direct experimental evidence (e.g. protein binding from yeast two-hybrid systems), and statistical models of high-throughput omics data (e.g. gene co-expression). Tools for network modelling of genetic data such as dmGWAS and EW_dmGWAS ^21^, DAPPLE ^22^ and “guilt by association” frameworks ^23^ are powerful extensions to the basic genetics toolbox. Furthermore, extensive tissue-specific network resources, such as the GIANT database ^24^, enable biologists to query the regulatory network context for genes of interest. Applications of network methods to multiple complex traits such as obesity ^25^, type 2 diabetes ^26, 27, 28^, coronary artery disease ^29,30^ and Alzheimer's disease ^19,31^ have led to successful identification of gene subnetworks (i.e. specific parts of the full network) of highly interconnected genes that represent pathogenic processes, and their central hubs or key drivers as potential points for intervention.

Despite the above methodological advances, several gaps remain to be addressed. The available methods are typically tailored for a particular combination of datasets (e.g. human genetics with gene expression, or human genetics with pathways or protein-protein interactions), thus lacking the flexibility to accommodate additional data types and multiple datasets from one or more species, tissues and platforms. Additionally, network approaches such as WGCNA ^32^ and postgwas ^33^ emphasize the detection of modules of co-operating genes, but validation experiments in the wet lab and therapeutic target selection require narrowing in on strong driver genes at the center of the module. Furthermore, the majority of the network tools start from a limited set of known top loci or genes and focus on ranking candidate genes based on network topology. One example is the GIANT database ^24^ and the NetWAS tool within, which provide convenient online query tools for such analyses. However, in most cases a full genomic analysis capable of extracting true subtle signals that are well below a significance cutoff from random noise is more powerful to achieve a comprehensive understanding of disease pathogenesis. Therefore, there remains a need for easily usable open source software that is designed for diverse types of genomic data to identify pathways, to model gene networks of diseases, and to pinpoint the key driver genes for further experiments in a streamlined and high-throughput manner.

To meet the challenge, we introduce Mergeomics, a flexible pipeline that integrates genomic associations, tissue-specific functional genomics resources, canonical pathways and weighted gene-gene interaction networks to identify pathogenic subnetworks and their key driver genes. The main components of Mergeomics are designed to 1) identify disease-associated subnetworks by aggregating genomic marker associations over functionally related or co-regulated genes; 2) perform pathway and network-level meta analysis across studies of different design, data type, platform, and species; 3) determine network key drivers by projecting disease subnetwork genes onto one or more system-scale gene or protein interaction networks. Here, we describe the methodology in detail and introduce new algorithms for pathway and network analyses. We also report the results from extensive testing of technical aspects such as parameter selection and data preprocessing based on simulated and empirical datasets to demonstrate that Mergeomics is statistically robust and outperforms previous methods. Finally, we applied Mergeomics to circulating cholesterol datasets, a clinically relevant human trait that is a major risk factor for cardiovascular disease. The most distinct aspects of Mergeomics lie in its applicability to both human and animal model studies, and adaptability for various types of association studies from GWAS, mutation burden from exome sequencing studies, and transcriptome-wide association studies (TWAS) to epigenome-wide association studies (EWAS) and metabolite or proteome-wide association studies. The source code for Mergeomics is released as an R package (http://mergeomics.research.idre.ucla.edu/Download/Package/).

## Results

### Overview of Mergeomics

Figure 1 shows the information flow within the Mergeomics pipeline. The Marker set enrichment analysis (MSEA), depicted on the left, combines disease association data (e.g., GWAS, EWAS, TWAS), functional genomics data from projects such as GTEx and ENCODE, and functional gene sets such as metabolic and signalling pathways and co-regulated gene modules. MSEA is based on the notion that while it is difficult to say which marker is causal for a disease, if the markers associated with a biological process (via their putative target genes selected based on functional evidence) are enriched for disease association signals, then it is plausible that at least some of those markers and their target genes are involved in causal disease mechanisms. The output from MSEA is a ranked list of gene sets that are significantly enriched for disease markers. We collectively denote these gene sets, which can be pathways, co-expression modules or gene signatures, as disease-associated gene sets. When multiple datasets of the same data type or different data types are available for a given disease or phenotype, the meta-MSEA component that is based on the same principles as MSEA but performs meta-analysis at the pathway or network level can be utilized.

**Figure 1.**
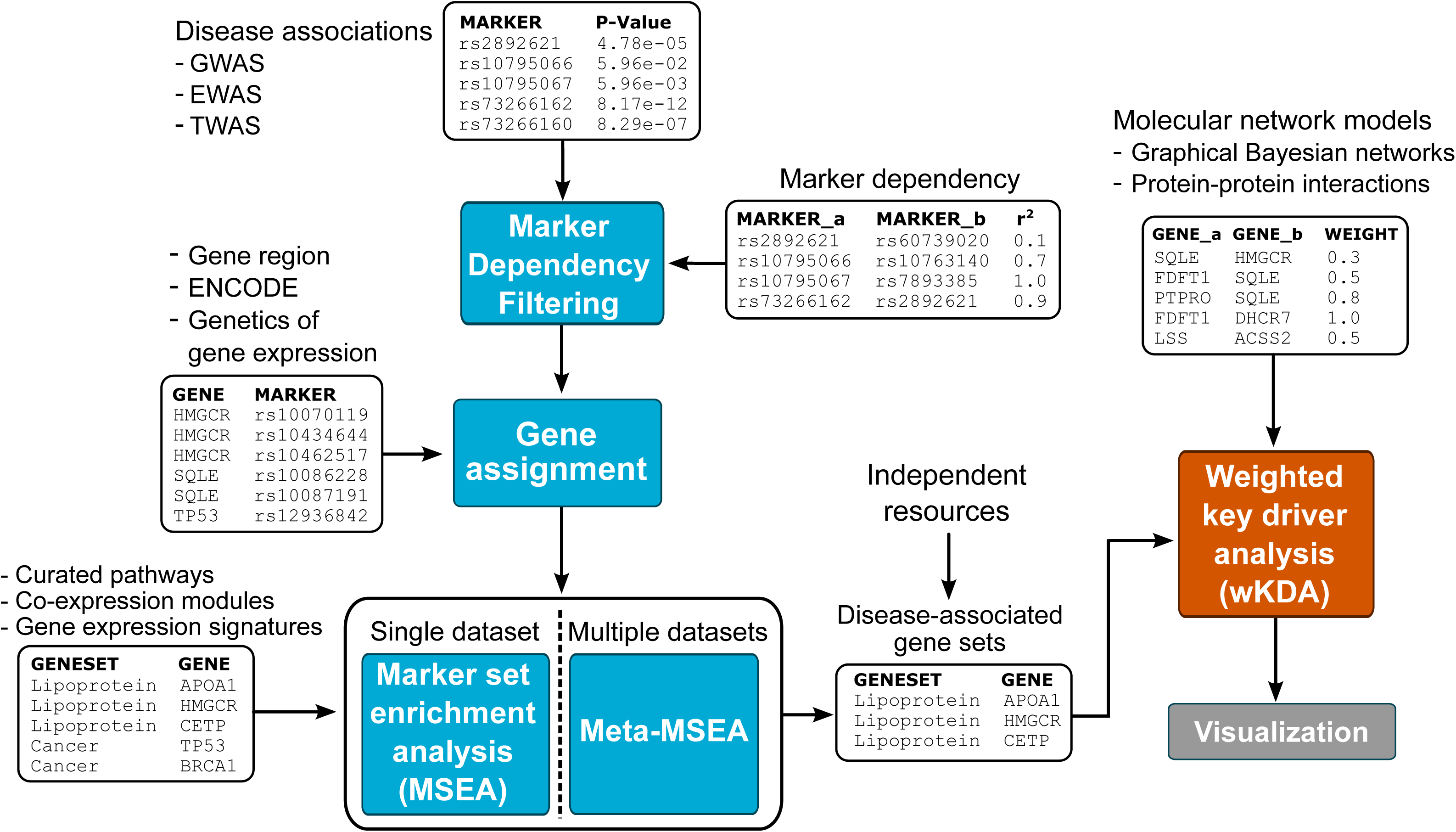
Input data formats and key analysis components of the Mergeomics pipeline.

The weighted Key Driver Analysis (wKDA, right side in Fig. 1) component of Mergeomics identifies local hubs that are central to the disease-associated gene sets by taking into consideration the network topology and the edge weight information between network nodes. This is accomplished by projecting the disease-associated gene sets from MSEA or meta-MSEA onto one or more types of gene or protein interaction networks containing detailed topology information, and then testing if the network neighbourhood of a particular hub shows over-representation of disease-associated genes. Hubs that demonstrate significant enrichment are defined as key drivers of the disease-associated gene sets.

Although MSEA (or meta-MESA) and wKDA are introduced as sequential steps in Mergeomics, they can be performed independently. MSEA or meta-MSEA can be performed without continuing to wKDA, and wKDA can be performed on pre-defined disease-associated genes without running MSEA or meta-MSEA.

### Calibration of MSEA

MSEA first converts a gene set from pre-defined functional pathways or co-regulated genes into a set of markers that are likely to perturb the function of the genes based on functional genomics data such as eQTLs and ENCODE information. The disease association P-values for this set of markers are then extracted from the summary statistics of a disease association study of interest. If there are a large number of small P-values in the marker set compared to what can be expected by chance, we conclude that the gene set we started from is enriched for disease associations. Key features of MSEA include: 1) it provides flexibility to accommodate association studies of different types or species, as long as the corresponding association statistics, marker-gene mapping, and pathway or gene set files are available; 2) it allows flexibility in gene-marker mapping to incorporate appropriate functional genomics information specific to the marker type (e.g., eQTL information between SNPs in GWAS and genes); 3) it allows marker filtering based on dependency measures between markers to select independent markers for statistical testing (e.g., linkage disequilibrium or LD information can be used to correct for linked SNPs in GWAS); 4) it utilizes a new test statistic with multiple quantile thresholds to automatically adapt to different association study datasets involving different sample sizes and statistical power; 5) it implements both marker-based and gene-based permutation strategies to estimate null distributions, with the latter adjusting for shared markers between genes and gene size.

To test the performance of MSEA, we performed simulation tests based on three cholesterol GWAS of varying sample sizes (a Finnish study of 8,330 individuals ^34^, the Framingham Heart Study with 7,572 participants ^35^, and Global Lipid Genetics Consortium or GLGC with 100,184 people ^36^) and a set of known causal lipid homeostasis genes from the Reactome pathway R-HSA-556833, “the metabolism of lipids and lipoproteins”. We resampled genes from this pathway into 100 sets of 25, 100 and 250 genes, respectively, to simulate 300 positive control signals of different magnitudes. Simultaneously, 300 corresponding sets of random gene sets were generated as negative controls. This procedure was repeated 100 times to produce stable statistics, and performance is evaluated as sensitivity, specificity and positive likelihood ratio (Details in methods).

We identified several important parameters that affect the performance of MSEA based on the three cholesterol GWAS datasets, including the percentage of top markers used, linkage disequilibrium cutoff to filter SNPs, and permutation type for null distribution estimation (**Supplementary Table 1**). First, the signal to noise ratio typically improved when genetic loci with relatively stronger associations rather than the full GWAS associations were used (**Supplementary Fig. 1**). This confirms previous findings for complex traits that heritability is maximally explained by the top portion of the GWAS SNPs ^37^. Second, the effect of LD correction depended on the permutation type, percentage of markers included, and the power of the association study under testing (**Supplementary Fig. 1**). In the marker-permuted MSEA, the marker labels were permuted to estimate the null distribution of the enrichment score for random expectation; in the gene-permuted MSEA, gene labels are permuted while the links between markers and genes are kept intact, thus more consistent with the hierarchical marker-gene-pathway cascade (**Supplementary Fig. 2**). In general, gene-based permutation is less sensitive to LD for more powered studies such as GLGC but performs better under higher LD cutoff for smaller studies such as Framingham; marker-based permutation is more sensitive to the percentage of markers used in MSEA particularly for smaller studies (**Supplementary Fig. 1**). In addition, the assignment of markers to their putative targets genes can be defined in multiple ways: chromosomal distance-based assignment based on the locations of gene regions, or functional information-based mapping determined by empirical data such as tissue-specific eQTLs or ENCODE information. Although empirical data is biologically more meaningful, to allow comparisons with other methods which mostly implement distance-based mapping, in our simulation analysis we used a window size of 20kb mapping between SNPs and genes by chromosomal location.

Overall, MSEA algorithm that relied on gene-based permutations demonstrated consistently high sensitivity, specificity and positive likelihood ratio with less parameter-dependent fluctuations compared to the marker-based version (**Supplementary Table 1**, **Supplementary Fig. 1**). Based on performance testing, we chose to use the top 50% of GWAS loci, an LD cutoff of r^2^ < 0.5, and gene permutation as the default setting. Of note, the differences due to datasets were typically larger than those due to parameters when using the gene-permuted MSEA (**Supplementary Fig. 3**).

### Performance comparison between MSEA, MAGENTA and i-GSEA4GWAS

MAGENTA ^15^ and i-GSEA4GWAS ^16^ are two widely used GWAS pathway analysis tools that are built upon an established gene set enrichment analysis ^38^. Both tools estimate the genetic associations for each gene, and then test if the aggregate gene score for a pathway is higher than expected. MAGENTA identifies the peak disease-associated SNP for each gene, and then adjusts the statistical significance of the peak SNP according to the size of the gene, LD and other potential confounders to produce the gene score. i-GSEA4GWAS uses a similar approach where a gene is considered significant if it contains any of the top 5% SNPs, and the pathway score is estimated by comparing the observed ratio of significant genes within the pathway against the expected ratio in the full set of genes that were covered by the GWAS. Compared to these methods, MSEA differs in test statistics, confounder adjustment, and flexibility in data accommodation.

The same simulated positive and negative control pathways that were used for calibrating MSEA were also used to compare the three different methods. Since this approach may give an unfair advantage to MSEA due to optimized calibration towards the positive controls, we also performed additional tests with 1,346 canonical pathways curated by Reactome ^39^, BioCarta (http://cgap.nci.nih.gov/Pathways/BioCarta_Pathways) and KEGG ^40^. All tests produced similar results: i-GSEA4GWAS lacked specificity and MAGENTA lacked sensitivity, whereas the gene-permuted MSEA provided the best balance and receiver operator characteristics. The results for the simulated positive and negative controls are depicted in Fig. 2, and the results from the additional tests with the canonical pathways are in **Supplementary Table 2**. Notably, the superior performance of MSEA over the other two established methods is more obvious when the GWAS involved smaller sample size and heterogeneous population.

**Figure 2.**
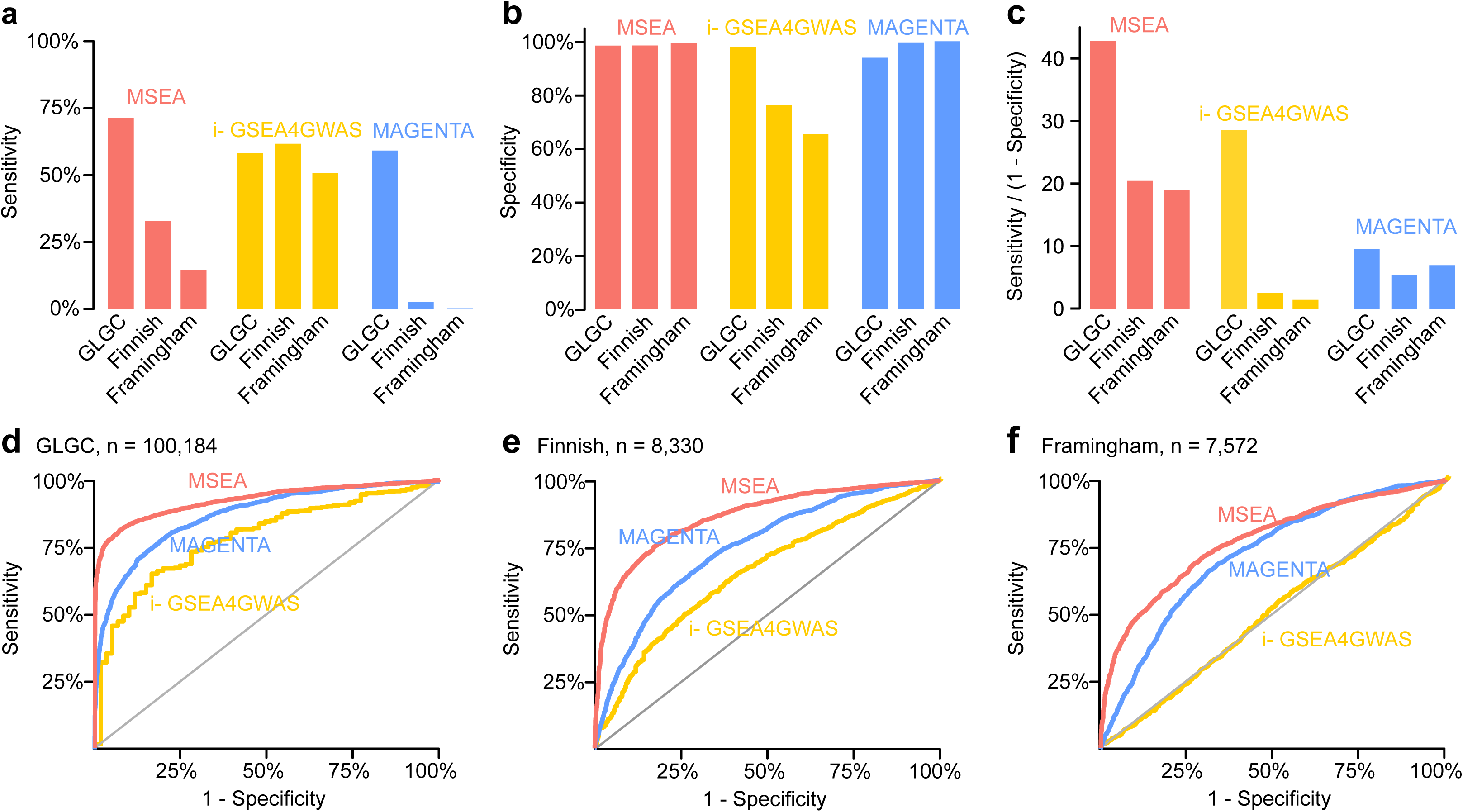
Comparison of three pathway enrichment methods across three GWAS. Only results for the gene-permuted version of marker set enrichment analysis (MSEA) are depicted (results for the marker-permuted version are in **Supplementary Table 1**). Performance is evaluated by sensitivity (**a**), specificity (**b**), positive likelihood ratio (sensitivity/(1-specificity)) (**c**) and receiver operating characteristic curve (**d-f**). Sensitivity was defined as the proportion of positive control pathways detected at FDR < 25%. Specificity was defined as the proportion of negative controls rejected at FDR > 25%.

### Meta-MSEA: pathway-level meta-analysis of multiple association studies

Various factors determine the quality of an association study. For example, GWAS result depends on the sample size, study design, accuracy of the phenotype, ethnicity and the coverage of the genotyping platform. Among the cholesterol GWAS we chose, the cholesterol level in the Finnish study was measured with a high-sensitivity NMR instrument in an ethnically homogeneous population, whereas the Framingham study relied on standard assays in a more mixed population. These differences may explain the lack of signals at FDR < 25% from the Framingham study in Fig. 2. However, the lipoprotein pathway was at the top of the list for all three GWAS, so a pathway-level meta-analysis can potentially boost weak but consistent signals. Importantly, unlike the traditional approach where meta-analysis is done at the marker level, this pathway-level analysis can bypass the need to match ethnicity or genotyping platforms, an advantage not present in the previous methods.

Mergeomics was specifically designed to produce output that is suitable for pathway-level meta-analysis (meta-MSEA) because pathway enrichment P-values are estimated from null distributions by parametric models (detailed in Methods). This ensures that the reported P-values are always greater than zero, and can be converted back to Z-scores by using the inverse Gaussian density function. i-GSEA4GWAS is an example where this procedure is difficult since only frequency-based P-values are estimated and highly significant signals can be set at P = 0. Table 1 lists the top pathways from meta-MSEA, and the full results for pathway-level meta-analysis are available in **Supplementary Data 1**. The pathway-level meta-MSEA not only accurately identified major lipoprotein and lipid transport pathways and the receptors that mediate lipid transfer to and from lipoprotein particles, but yielded more significant P-values than those obtained from the pathway analysis of conventional meta-GWAS conducted at the SNP-level. Such superiority of meta-MSEA was consistently observed using simulated gene sets (**Supplementary Fig. 4**). These results demonstrate the pathway-level meta-analysis is more powerful than the traditional SNP-centric approach to meta-analysis when investigating the genetic perturbations to biological processes, and we have accordingly incorporated the support for multiple association studies in the Mergeomics pipeline. Importantly, this pathway level meta-analysis feature allows integration of different types of omic association data sets. For example, association studies for a particular disease done at genetic, gene expression, epigenetic, and metabolite levels can be meta-analyzed after conducting MSEA on each association dataset, allowing the detection of functional pathways or networks that are perturbed by different types of molecular entities.

**Table 1.** Top 15 pathways associated with cholesterol levels in three GWAS datasets among 1,346 canonical pathways tested. Values in the cells represent -logi_0_-transformed enrichment P-values from MSEA of individual cohorts and meta-MSEA. MSEA was run with gene permutation, top 50% of markers and LD cutoff r^2^ < 50%. Only pathways with meta-MSEA -log_10_P > 4.0 were included here; the full results are available in **Supplementary Data 1**. The column 'Meta-GWAS' was produced by first conducting SNP-level meta-analysis using the SNP associations from all three GWAS by inverse-variance meta-analysis and then estimating the pathway enrichment. The column 'Meta-MSEA' was produced by pathway-level meta-analysis in which pathway enrichment Z-scores from each GWAS were first estimated with MSEA, followed by addition of these approximate Z-scores into a single combined enrichment signal before calculating the statistical significance. The adjusted 5% Bonferroni significance level for 1,346 independent tests is at –log_10_P > 4.43.

| Pathway | FIMM | Framingham | GLGC | Meta-GWAS | Meta-MSEA |
| --- | --- | --- | --- | --- | --- |
| Lipid digestion, mobilization, and transport | 4.2 | 5.5 | 6.1 | 5.0 | **13.8** |
| Lipoprotein metabolism | 4.7 | 4.8 | 5.9 | 5.4 | **13.5** |
| Chylomicron-mediated lipid transport | 4.9 | 4.9 | 4.7 | 5.0 | **12.6** |
| Metabolism of lipids and lipoproteins | 3.2 | 1.7 | 6.2 | 3.6 | **8.5** |
| Cytosolic tRNA aminoacylation | 3.6 | 2.1 | 1.9 | 2.7 | **5.9** |
| Binding and Uptake of Ligands by Scavenger Receptors | 1.9 | 2.3 | 3.4 | 2.9 | **5.9** |
| Scavenging by Class A Receptors | 1.8 | 2.2 | 3.2 | 3.5 | **5.6** |
| Metabolism | 1.8 | 1.5 | 3.9 | 3.0 | **5.4** |
| PPARA Activates Gene Expression | 1.7 | 2.2 | 2.8 | 1.3 | **5.1** |
| Retinoid metabolism and transport | 1.0 | 2.8 | 3.0 | 1.4 | **4.9** |
| Regulation of Lipid Metabolism by | 1.3 | 2.0 | 2.8 | 1.6 | **4.5** |
| Peroxisome proliferator-activated<br>receptor alpha (PPARalpha) |  |  |  |  |  |
| Fatty acid, triacylglycerol, and ketone<br>body metabolism | 1.5 | 1.7 | 2.5 | 1.6 | **4.1** |
| Clathrin derived vesicle budding | 1.9 | 1.3 | 2.4 | 1.3 | **4.0** |
| Diseases associated with visual<br>transduction | 1.4 | 1.9 | 2.2 | 2.3 | **4.0** |
| ABC transporters | 1.8 | 0.9 | 3.2 | 2.8 | **4.0** |

### Weighted key driver analysis (wKDA) to detect disease regulators

The MSEA or meta-MSEA component of Mergeomics identifies pathways or co-regulated gene sets that are perturbed in a disease. However, the interactions between genes within these disease-associated gene sets are not evident. To this end, a key driver analysis (KDA) was previously developed to detect important hub genes, or key drivers, whose network neighbourhoods are over-represented with disease associated genes ^29,41^. The key driver concept is based on the projection of the disease-associated gene sets onto a separate graphical network model of gene regulation that represents molecular interactions in the full system (Fig. 3a). However, KDA ignores the edge weight information generated by most network inference algorithms. As edge weight typically represents association strength or reliability of the connection between genes, this data carries valuable topological information. Here, we introduce a new algorithm wKDA that takes into account edge weights to increase accuracy, and a new co-hub concept that considers the local topology around hub genes to reduce multiple testing (Fig. 3). wKDA starts by searching a graphical network for candidate hub genes and ignores genes with few connections. It then collects the neighbouring genes for each candidate hub, and estimates the contribution of the disease-associated genes within the neighbourhood of the hub. If the contribution is stronger than what would be expected by chance, we conclude that the hub is a key driver of the disease-associated gene sets. Moreover, we developed a co-hub concept for wKDA (Fig. 3b), given the fact that networks with densely connected communities may complicate the interpretation of the key driver signals. Basically, if a subnetwork of genes has multiple highly interconnected genes at the center, it is critical to consider them collectively. For practical purposes, we select one of the central genes as the independent hub, and the rest as co-hubs. The rationale is two-fold: first, the statistical power is increased by only considering the independent hubs when adjusting P-values, as they also capture the signals from their respective co-hubs. Second, the co-hub concept is a useful qualitative measure when selecting the most promising subnetworks and key drivers for experimental validation. For instance, if a key driver has co-hubs with known functions, these can give clues as to the role of poorly understood genes. On the other hand, if a key driver is to be perturbed in an experiment, it may be important to incorporate the co-hubs as integral parts of the experimental design.

**Figure 3.**
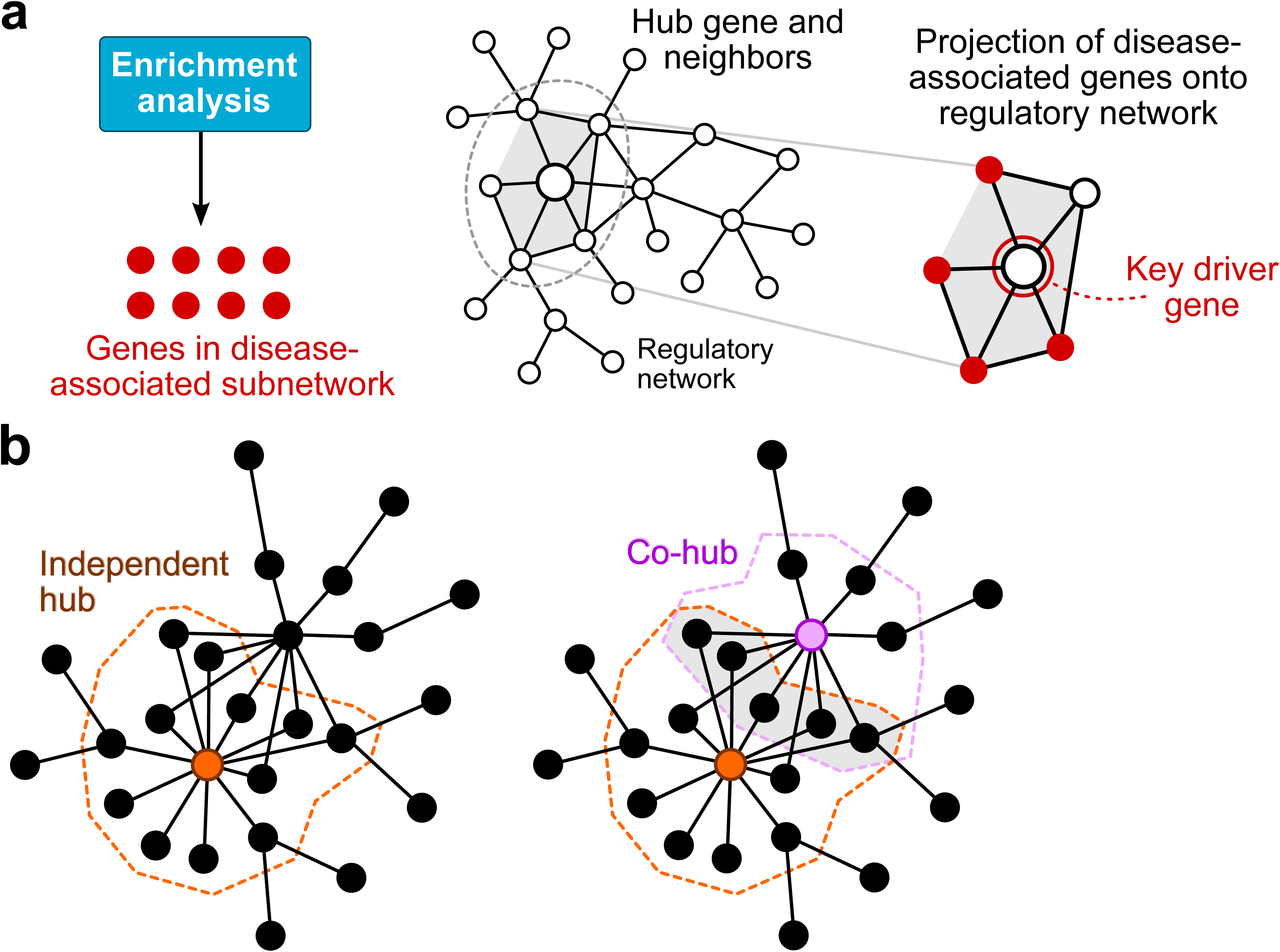
Schematic illustration of the concept of a key driver gene (**a**) and local hubs with overlapping neighborhoods (**b**).

The main difference between KDA and wKDA involves the counting of the subnetwork members around the hub. In KDA, each node is treated equally without consideration of the edges, and the enrichment is based on the excess proportion of disease genes within the hub neighbourhood. The new wKDA assigns a larger weighting coefficient for a node with higher edge weights than for other nodes in a network neighbourhood (details in Methods). Therefore, if the disease genes have higher edge weights to a hub or its neighbors, the enrichment score will be higher. From a practical perspective, the previous KDA detects key drivers that are connected to a large proportion of the disease-associated subnetwork genes, whereas the wKDA tends to detect key drivers that have high-weight edges to disease genes. In addition to identifying key drivers and co-hubs, wKDA also outputs Cytoscape input files for the key drivers and their local subnetworks with disease genes highlighted that could be visualized in the Cytoscape software ^42^.

### Performance of wKDA and comparison with KDA

To evaluate the performance of wKDA in comparison to the unweighted KDA, we first set up three disease-associated gene sets to test against four gene regulatory networks. The three gene sets included two lipid subnetworks (denoted as Lipid I and Lipid II) derived from our previous study ^29^ and the R-HSA-556833 (Metabolism of lipids and lipoproteins) pathway from Reactome. The gene-gene interaction network models were probabilistic Bayesian gene regulatory networks constructed from multiple adipose and liver datasets (**Supplementary Table 3**). We organized these networks into two independent weighted adipose networks and two independent weighted liver networks using non-overlapping datasets, where edge weight represents the estimated reliability of an edge between genes.

We used the overlap ratio (defined in Methods) of the identified key driver genes between the two independent networks of the same tissue to assess the predication accuracy of wKDA and KDA. As shown in Fig. 4, the new wKDA outperformed the previous unweighted KDA for all three gene sets against independent networks in both tissues. To test the sensitivity of the key driver approach, we also partially randomized the adipose and liver networks as a model of topological noise. As expected, when some of the edges were randomly rewired, the number of consistent key drivers between two independent networks of the same tissue declined, and when all edges were rewired, no consistent key drivers were detected (Fig. 4). Notably, wKDA was able to detect consistent signals even when half the network was rewired, thus demonstrating the inherent robustness of the wKDA concept compared to the unweighted version. Importantly, because wKDA was specifically designed for weighted networks whereas the unweighted KDA mainly focuses on the network topology without considering weight information, key drivers with high-weight (i.e., high reliability) edges between subnetwork genes were preferred by wKDA. This difference likely explains the better reproducibility of wKDA compared to the unweighted KDA (Fig. 4, **Supplementary Table 4**).

**Figure 4.**
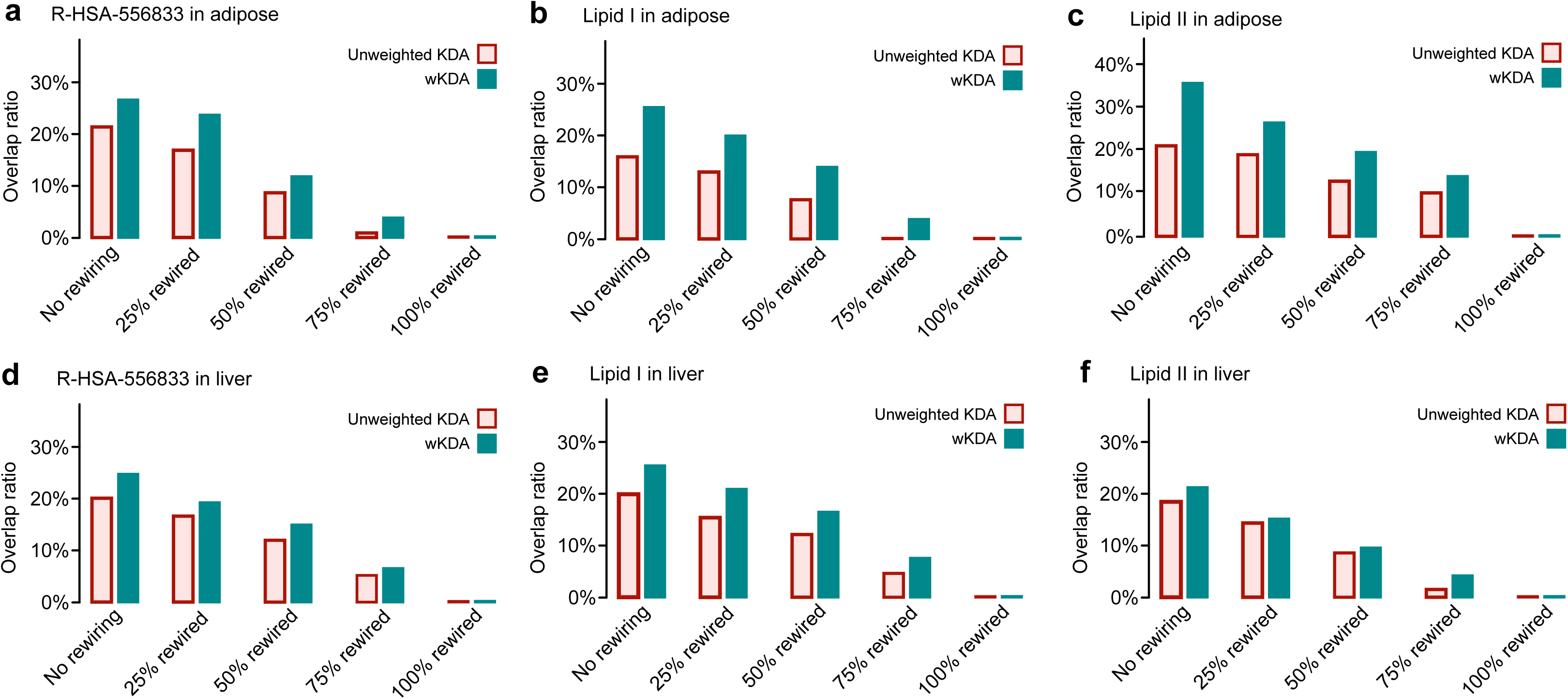
Performance comparison between the original unweighted key driver analysis, and the new algorithm designed for weighted networks (wKDA). Two empirical subnetworks (Lipid I & II) were obtained from a previous publication ^29^, and a canonical metabolism of lipids and lipoproteins pathway was obtained from the Reactome database (R-HSA-556833). The methods were tested by projecting the three functional subnetworks onto two independent adipose networks (**a-c**) and two independent liver regulatory networks (**d-f**). The adipose and liver networks were constructed from a collection of Bayesian tissue-specific network models (**Supplementary Table 3**). Overlap between the tissue-specific key driver signals across two independent regulatory networks was defined according to the formula in Methods. Overlap ratio was calculated for both original networks and networks rewired at 25%, 50%, 75%, 100%.

### Case study: application of Mergeomics to circulating cholesterol datasets

The cholesterol in low-density lipoprotein particles has been established as a causal biomarker for cardiovascular disease ^43^. Multiple large GWAS have revealed a complex genetic regulation of circulating cholesterol that is likely to involve multiple genes ^36^, making cholesterol genetics an interesting test case for our pipeline. Furthermore, the decades of research have catalogued the features of cholesterol biosynthesis and lipoprotein transport at the pathway level, which makes it easier to verify that our methodology produces meaningful biological results.

Significant pathways related to total cholesterol are already identified in the aforementioned MSEA and meta-MSEA analyses (**Supplementary Data 1**). Table 1 lists the top 15 pathways that were genetically perturbed by cholesterol-associated loci based on the ranks in the meta-MSEA results. Aside from the expected hits for lipoprotein transport, several pathways related to cellular lipid trafficking (scavenging receptors and ATP-binding cassette transporters of class A) and lipid metabolism (such as fatty acid, triacylglycerol and ketone body metabolism) were identified. Interestingly, the top hits included ‘Cytosolic tRNA aminoacylation’ and 'PPAR-alpha activates gene expression', which suggest that these transcriptional regulatory processes are intrinsically intertwined with the traditional concepts of enzyme-driven metabolic pathways in cholesterol biosynthesis and transport.

Because of the presence of overlaps in gene memberships between certain curated pathways, we merged 82 overlapping pathways with meta-MSEA p-value < 0.05 into 43 non-overlapping gene “subnetworks” at a maximum allowed overlap ratio of 0.20, and performed a second run of meta-MSEA using these merged subnetworks to retrieve the top six subnetworks (**Supplementary Table 5**). The strongest signal was observed for Subnetwork 1 (P < 10^−16^) that contained genes encoding key apolipoproteins and lipid transport proteins (such as *LDLR, CETP* and *PLTP)*. Subnetwork 2 (P < 10^-8^) included genes related to lipid biosynthesis and catabolism (including the statin target *HMGCR)*, oxidoreductive enzymes, metalloproteins and mitochondria. Subnetwork 3 represents a biologically intriguing connection between circulating cholesterol and the immune system: it contained proteins that are involved in the transport of fatty acids and lipids in blood (Albumin and apolipoproteins A1, B, A and L1), collagen genes, and the immunoglobulin family. Subnetwork 4 mainly contained the ATP-binding cassette family of transmembrane transporters responsible for lipid and cholesterol transfer across cell membranes. Subnetwork 5 included genes for metabolizing retinoid, an important mediator of cholesterol transport. Subnetwork 6 connected transcriptional regulation with fatty acid metabolism.

Using wKDA, we identified candidate key drivers in the liver and adipose tissues for each of the top six cholesterol-associated subnetworks (top five along with co-hubs listed in Table 2 and full list in **Supplementary Data 2**). As exemplified in Fig. 5a, the top adipose key driver for Subnetwork 2 is the very long chain acyl-CoA dehydrogenase *(ACADVL)*, which catalyzes the first step in mitochondrial beta-oxidation. Notably, the two co-hubs for *ACADVL (PPARA* and *CIDEA*) are also highly relevant genes for maintaining lipid homeostasis: *PPARA* is one of the master regulators of lipid metabolism with clinically approved class of drugs (fibrates) already in use; *CIDEA* has been linked to apoptosis, and mouse knock-outs have demonstrated significant effects on the metabolic rate and lipolysis ^44^. In liver (Fig. 5b), the top key driver of Subnetwork 2 is fatty acid synthase *(FASN)*, which is a key driver in adipose tissue as well. The second top key driver squalene epoxidase *(SQLE)* and its co-hubs *(FDFT1, IDI1, MSMO1, NSDHL, HMGCS1, ALDOC)* either catalyze or regulate cholesterol biosynthesis. *HMGCR*, although not listed as top five key drivers, is a highly significant key driver (*P* < 10^−14^) and a co-hub of *MMT00007490*. Subnetwork 2 and Subnetwork 6 shared multiple common key drivers in the adipose network (Fig. 5a). These included aconitase 2 *(ACO2)*, an enzyme that catalyzes citrate to isocitrate in mitochondrion, and *ACADVL* and its co-hubs. Perturbation of the majority of the top key drivers, including *ACADVL, FASN, SCD, ACO2, COL1A2, POSTN, EHHADH, DHCR7, HSD17B7, GC, AQP8, INSIG1*, has been shown to cause abnormal cholesterol and lipid homeostasis based on the Mouse Gene Informatics database and International Mouse Phenotyping Consortium ^45, 46^. In summary, biological plausibility, literature evidence and experimental data all support the fundamental role of the top key drivers in regulating cholesterol metabolism.

**Figure 5.**
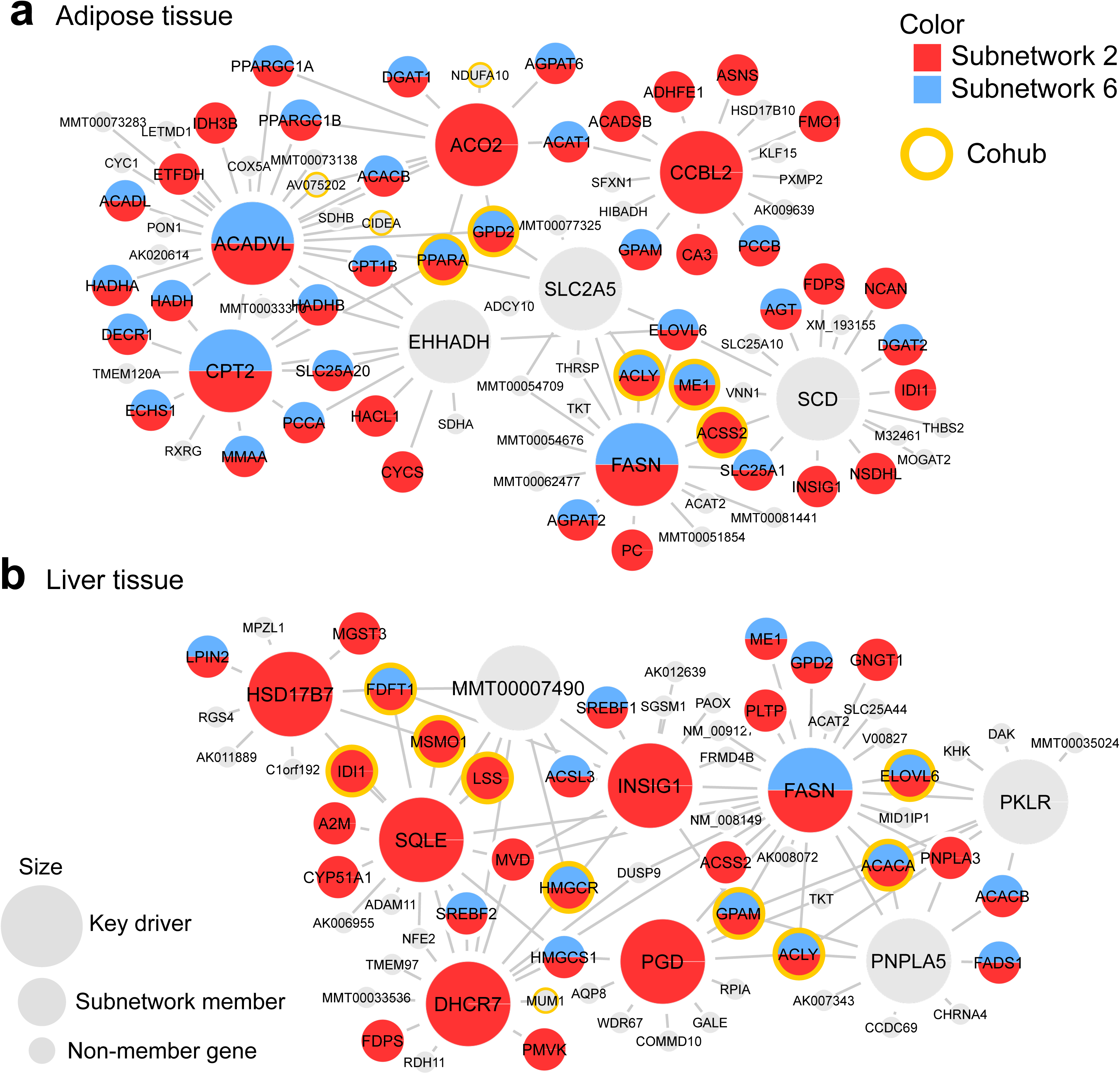
Visualization of adipose (**a**) and liver (**b**) networks around top key drivers identified for cholesterol-associated subnetworks. Top key drivers (Nodes with the largest size) are selected as the top five independent key regulatory genes (genes whose neighbourhood has less than 25% overlap with the neighbourhood of other independent hubs) for subnetwork 2 and subnetwork 6. Subnetwork member genes are denoted as medium size nodes and non-member genes as small size nodes. Top co-hubs (co-hubs with FDR<1e-10 in wKDA) are also highlighted by yellow circle. Only edges that were supported by at least two studies were included.

**Table 2.**
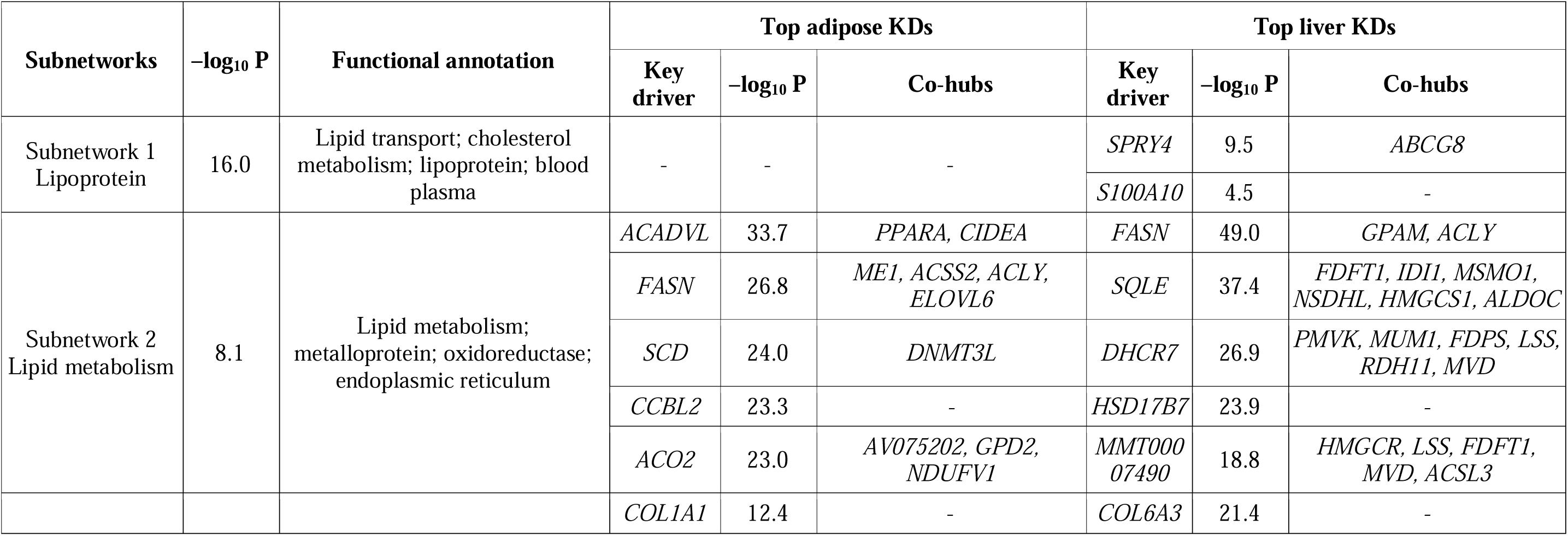
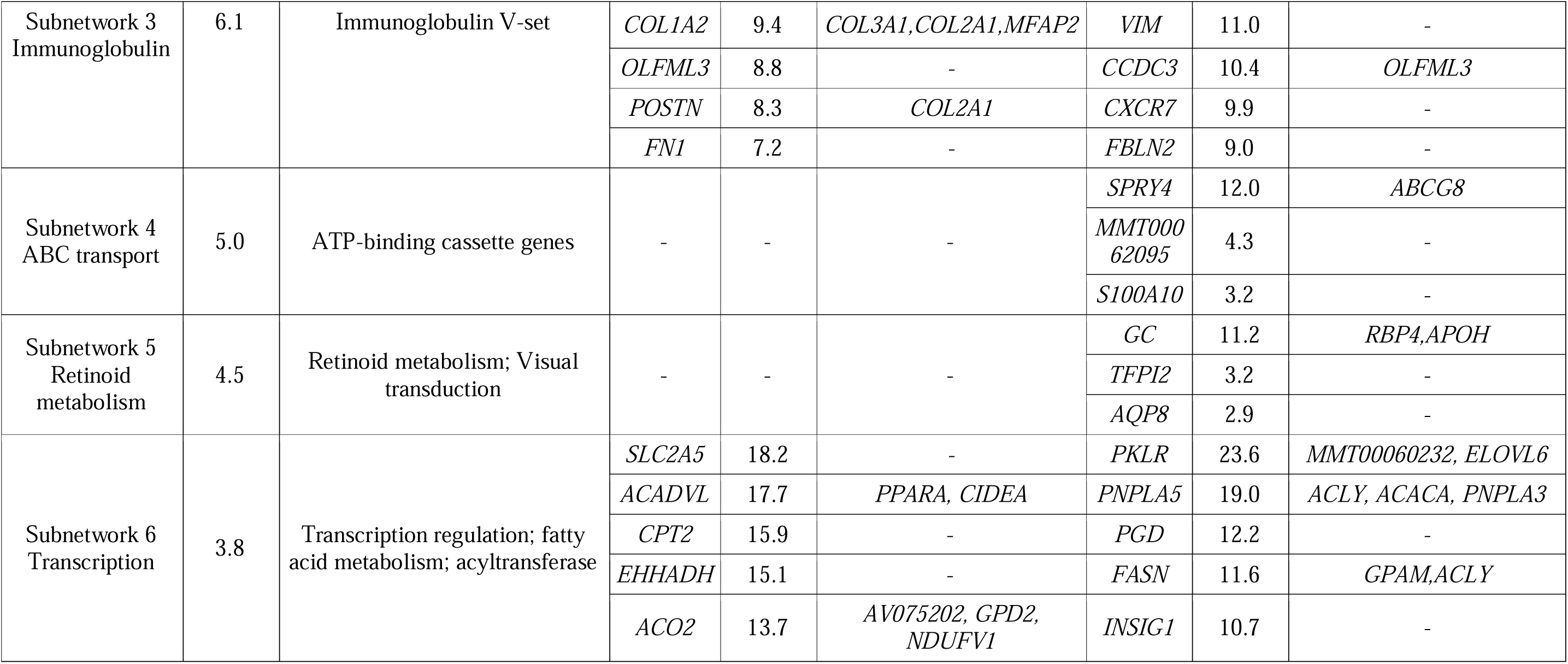
Cholesterol subnetworks after merging cholesterol-associated canonical pathways and their respective top five key drivers (FDR<1%). To confirm that the merged subnetworks were genetically associated with cholesterol, a second round of MSEA was performed. Functional annotations were determined with the DAVID Bioinformatics Tool ^52^. Statistical significance was estimated as described in Table 1. The adjusted 5% Bonferroni significance level for 43 independent merged supersets is at –log_10_P > 2.93. For key drivers, the P-value indicates the enrichment of cholesterol-associated genes around a local hub in a tissue-specific network. Before analyses, Bayesian networks from multiple studies were combined to create weighted adipose and liver consensus networks. Gene symbols have been translated to human when available. Co-hubs were defined based on the overlap of neighboring genes in the network: if the overlap ratio of two local hubs was above 33% (50% of the combined neighborhood was shared between the hubs), the hub with the lower P-value for the subnetwork enrichment was chosen as the key driver, and the other hub was designated as a co-hub.

## Discussion

The explosion of genomics data provides unprecedented opportunities to identify important mechanisms of disease across studies. Here we introduce a standardized pipeline to connect disease association studies with functional data and curated knowledge, and apply it to the genetics of cholesterol. We used multiple independent datasets to show how the Mergeomics components (MSEA and wKDA) outperformed previous methods in sensitivity and specificity. The generic nature of MSEA makes it straightforward to apply it to individual genomic association studies in different species and different omics data types. The unique pathway-level meta-analysis feature makes it highly powerful in overcoming population and study design differences to integrate diverse data sources to accurately identify shared biological processes across studies. The weighted network algorithm for wKDA is equally flexible: it can be applied to diverse biological networks and provides statistical and qualitative information on the key regulating genes in a tissue- and network type-specific fashion. Our case study suggested that there is an underlying gene regulation pattern that involves existing drug targets (such as *PPARA* and *HMGCR*) as well as less known genes (such as *ACADVL* and collagen genes), that can help explain the complex signals from cholesterol genetic studies, and guide the development of novel hypotheses and wet lab experiments. With the release of the R library, we provide the scientific community with easy-to-use tools to make sense of the exiting mass of genomics resources.

## Methods

### Market set enrichment analysis

The default setting of MSEA takes as input 1) summary statistics from genomic association studies, 2) measurement of relatedness or dependency between genomic markers, 3) functional mapping between markers and genes, and 4) functionally defined gene sets (e.g., biological pathways or co-regulated genes). For GWAS, SNPs are first filtered based on the LD structure to select for only SNPs that are relatively independent given an LD threshold ^29^. For other types of association studies, correlations between co-localized markers may be used. For a given gene set, gene members are first mapped to markers based on the functional mapping file and then the disease association p values of the corresponding markers are extracted to test for enrichment of association signals. To test enrichment, both a gene-based analysis and marker-based analysis are implemented in MSEA.

The null hypothesis for the enrichment of association signals within a gene set can be defined as

> *Gene-based H_0_: Given the set of all distinct markers from a set of N genes, these markers contain an equal proportion of positive association study findings when compared to all the distinct markers from a set of N random genes*

or as

> *Marker-based H_0_: Given a set of M distinct markers, these markers contain an equal proportion of positive association study findings when compared to a set of M random markers.*

We only focus on distinct markers to reduce the effect of shared markers among gene families that are both close in the genome and belong to the same pathway (and presumably have overlapping functionality). Furthermore, our software has a feature that merges genes with shared markers before analysis to further reduce artifacts from shared markers. The expected distribution of the test statistic under the null hypothesis can be estimated empirically by randomly shuffling the gene or marker labels (**Supplementary Fig. 2**). The gene-based approach is robust against LD and other artifacts. The marker-based approach is more sensitive, however it requires substantial correction for LD to be reliable and may suffer from artifacts due to the non-random positional patterns of gene regions in the genome.

To avoid assessing enrichment based on any pre-defined association study P-value threshold (e.g., p<0.05) which can mean different association strengthes in studies of varying sample size and power, we developed a new test statistic with multiple quantile thresholds to automatically adapt to any dataset:

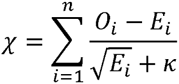

In the formula, *n* denotes the number of quantile points, *O* and *E* denote the observed and expected counts of positive findings (i.e. signals above the quantile point), and κ = 1 is a stability parameter to reduce artefacts from low expected counts for small gene sets. The frequency of permuted signals that exceed the observation is determined as the enrichment P-value. For highly significant signals where the frequency-based value is zero (i.e. no permuted signal exceeds the observation), we fit a parametric model to the simulated null distribution to approximate the corresponding Z-score (see details under the section “adaptive Gaussian approximation for estimating P-values” below). For meta-MSEA of multiple association studies, pathway enrichment Z-scores from each dataset are first estimated with MSEA, followed by addition of these approximate Z-scores into a single combined enrichment score for P value estimation.

### MSEA performance evaluation

The MSEA within Mergeomics can be reconfigured depending on the type of dataset and study design. We identified several parameters that could affect the performance of the pipeline such as marker filtering by including top associated markers based on a percentage cutoff, dependency or relatedness (such as LD) cutoff for pruning redundant markers, and the mapping between genes and markers. Here we focus on the marker filtering percentage and LD cutoff as they represent the two key technical challenges. Of note, the mapping between genes and markers can be defined empirically ^11, 13^, but we used a chromosomal distance-based approach for testing to make Mergeomics consistent with most of the current pathway enrichment tools. In fact, for GWAS, the assignment of SNPs to their target genes based on their chromosomal location is the commonly adopted approach in other methods, whereas Mergeomics allows users to apply any available assignment method, including the data from tissue-specific eQTL studies and ENCODE.

High cholesterol is a major risk factor for cardiovascular disease, and cholesterol metabolism and transport is one of the most studied and understood areas of human biology, which makes cholesterol GWAS ^36^ an informative dataset for method assessment. GWAS summary data for circulating cholesterol were available from 7,572 individuals in the Framingham Study ^35^, 8,330 Finnish individuals ^34^, and 100,184 participants from the Global Lipid Genetics Consortium ^36^. The GLGC contains the two smaller studies, but as the total overlap was less than 10% between the datasets, we assume that the three GWAS are independent for the purposes of this study. All participants were predominantly Caucasian descent, and we used the corresponding LD data from HapMap ^47^ and 1000 genomes project ^48^ in our analyses.

We simulated true positives and true negatives to determine a suitable combination of parameters and to compare performance of different methods. We collected genes from the Reactome pathway R-HSA-556833, “the metabolism of lipids and lipoproteins”, treating these genes as true signals related to cholesterol and lipid metabolism. These genes were randomly grouped into 300 positive control pathways, including 100 with size 25, 100 with size 100, 100 with size 250, respectively. Simultaneously, 300 negative control pathways with the same size distribution as the positive control pathways were generated by randomly selecting genes from the non-cholesterol gene pool consists of 8633 genes from the pathway database of Reactome ^39^, BioCarta (http://cgap.nci.nih.gov/Pathways/BioCarta_Pathways) and KEGG ^40^. These manually generated pathways were combined with 1,346 original canonical pathways and analysed by MSEA, MAGENTA and i-GSEA4GWAS. The performance was evaluated as sensitivity (number of positive control pathways at FDR < 25% divided by total number of positive control pathways), specificity (number of negative control pathways at FDR < 25% divided by total number of negative control pathways) and the likelihood to pick up true positive pathways (Positive Likelihood Ratio), calculated as sensitivity / (1 - specificity).

### wKDA

**Supplementary Fig. 5** depicts the main components of wKDA which utilizes both the network topology information and the edge weight information when available. In wKDA, the network topology is first screened for suitable hub genes whose degree (number of genes connected to the hub) is in the top 25% of all networks nodes (**Supplementary Fig. 5**, middle box on the left). We further classify these genes as either independent hubs or co-hubs, where co-hub is defined as a gene that shares a large proportion of its neighbours with an independent hub. First, the candidate independent hubs are sorted according to the node degree, from low to high. This is to ensure that we capture local structures rather than one master hub that covers the majority of the network (e.g. housekeeping genes would make poor drug targets due to global side-effects). Next, the sorted hubs are tested one by one for neighbourhood overlaps with the already accepted hubs. If sufficient overlap (as defined under section “definition of overlap between two gene sets” below, default value is 33%) is detected, the current hub is assigned as a co-hub for the previously accepted overlapping hub.

### wKDA statistics

Once the hubs and co-hubs have been defined, the disease-associated gene sets that were discovered by the MSEA are overlaid onto the network topology to see if a particular part of the network is enriched for the potential disease genes. First, the edges that connect a hub to its neighbours are simplified into node strengths (strength = sum of adjacent edge weights) within the neighbourhood (**Supplementary Fig. 5**, **Plots B-D**), except for the hub itself. For example, the top-most node in Plot C has three edges that connect it with the other neighbors with weights that add up to 7 in Plot D. By definition, the hub at the center will have a high strength which will skew the results, so we use the average strength over the neighbourhood for the hub itself. The reduction of the hub neighbourhood into locally defined node strengths improves the speed of the algorithm and makes it easier to define an enrichment statistic that takes into account the local interconnectivity. In particular, the weighting of the statistic with the node strengths emphasizes signals that involve locally important genes over isolated peripheral nodes. In Plot D of **Supplementary Fig. 5**, the overlap between the hub neighbourhood and a hypothetical disease-associated gene set is indicated by the circles around the top three nodes. The sum of the respective strengths is 15, which represents 57% of the total sum of 26.4 in the neighbourhood (pie chart in Plot D). The final enrichment score is estimated with the formula below.

The null hypothesis for the enrichment of disease genes within a subnetwork can be expressed as

> *Weighted key driver H_0_: Given the set of nodes adjacent to a key driver, and with each node having a local strength as estimated by their mutual connectivity, the ratio of disease gene-member sum of strengths to the total sum of strengths is equal to the ratio for a randomly selected gene set that matches the number of disease genes.*

The test statisitic for the wKDA is analogous to the one used for MSEA

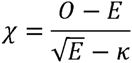

except that the values *O* and *E* represent the observed and expected ratios of pathway memberships. In particular,

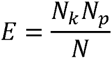

is estimated based on the hub degree *N_k_*, pathway size *N_p_* and the order of the full network *N*, with the implicit assumption that the weight distribution is isotropic across the network.

Statistical significance of the disease-enriched hubs, henceforth key drivers, is estimated by repeatedly permuting the gene labels and estimating the P-value based on the simulated null distribution. To control for multiple testing, we perform adjustments in two tiers. First, the P-values for a single subnetwork are multiplied by the number of independent hubs (Bonferroni adjustment). All hubs with adjusted P > 1 are discarded. For random data, the truncated results will be uniformly distributed between 0 and 1, and hence they can be treated as regular P-values. In the second stage, all the P-values for the subnetworks are pooled and the final false discovery rates are estimated by the Benjamini-Hochberg method ^49^.

### Performance assessment of wKDA

In this study, we use the Bayesian networks ^50, 51^ constructed from published genomic studies where both DNA and RNA are collected from adipose and liver tissue samples (**Supplementary Table 3**). Given that SNPs affect gene expression but not vice versa, it is possible to infer the causality of regulation between two correlated genes by looking at the statistical associations between SNPs and gene transcripts. For instance, if only one of two co-expressed genes are regulated by a SNP, it is possible that the SNP regulates gene A expression that in turn regulates gene B expression, thus showing a causal link between gene A and B. However, if the genes are each regulated by the same SNP, it may be the sign of an incidental co-expression without a direct causal relationship. In a Bayesian model, the uncertainty over the causality is estimated by conditional probabilities between co-expressed genes, and the structure of the resulting network is further constrained to an acyclic topology to ensure computational feasibility ^50, 51^.

We organized the individual Bayesian networks into two independent weighted adipose networks and two independent weighted liver networks from non-overlapping datasets (**Supplementary Table 3**), where edge weight represents the estimated reliability of a connection or edge between genes based on the consistency of the edge between datasets. Using these networks and three test gene sets related to lipid metabolism as inputs, we ran wKDA and the previously developed unweighted KDA to identify liver and adipose key drivers of the lipid gene sets. To assess the prediction accuracy of wKDA and KDA, we used the overlap ratio of the identified key driver genes (as defined under section “definition of overlap between two gene sets” below) between the two independent networks of the same tissue.

### Adaptive Gaussian approximation for estimating P-values in MSEA and wKDA

The exact shape of the null distribution is dependent on the size of the gene set and on the mapping between the genes and the markers (MSEA) or on the size and topology of the gene network (wKDA). To estimate the P-value from these various permutation approaches, we created a generic algorithm for a parametric approximation using the Gaussian function. In the range where a direct frequency-based P-value is accurate (i.e. with 10,000 permutations it is possible to accurately estimate P-values above ~0.001), we found that the Gaussian approximation was highly concordant. For P < 0.001, we found that the Gaussian model produced biologically plausible rankings of statistical significance. We tested other models, but found that the potential benefit from using more long-tailed distributions was outweighed by the difficulties in applying them in practice. For instance, the t-distribution was more conservative than the Gaussian estimate, but assigning an appropriate degree of freedom was problematic given the diverse nature of the null hypotheses.

Let X denote the series of simulated test statistics (as defined in the previous section) from the permutation analysis. Then the transformation algorithm can be expressed as

1) α = min(X_0_), X_1_ = X_0_ - α
2) β = median(X_1_), X_2_ = X_1_ / β
3) X_3_ = log(γX_2_ + 1)
4) μ = mean(X_3_), σ = sd(X_3_)
5) Evaluate how well X_3_ approximates N(μ, σ)
6) If necessary, try a different γ and go back to Step 3.

The parameters from Steps 1–4 can be saved and reapplied to new data, which makes it possible to determine the transformation exclusively based on simulated statistics, and then apply it to the observed test statistic to yield the parametric enrichment score

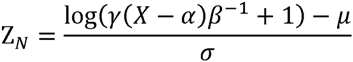

The rationale for Gaussian approximation is based on the attractive analytical properties of Gaussian distributions. Nevertheless, if the approximation is inaccurate, the results can be biased and lead to erroneous conclusion. In particular, any dependencies between markers tend to elongate the tails of the “true” distribution when using marker permutations for the MSEA. For this reason, we also report the raw frequency of false positive findings from the permutation analysis for each gene set.

### Definition of overlap between two sets

We define the overlap between two sets *A* and *B* as

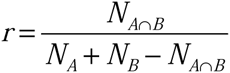

where *N* denotes the number of items. The ratio is zero when there are no shared genes and one when *A = B*. Importantly, the ratio is symmetric for two sets of different sizes i.e. the labels A and B can be swapped without affecting the value of *r*. This definition is used for the calculation of overlap ratio when merging overlapping significant gene sets from MSEA, determining hub-cohub relationship, as well as evaluating the consistency of key drivers identified in independent Bayesian networks.

### Availability

Mergeomics is available as a freely downloadable R package (http://mergeomics.research.idre.ucla.edu/Download/Package/). The package supports full Mergeomics functionality, plus the option to generate Cytoscape input files for quick network visualization. At the download site, sample omics datasets, network models, and a standalone C++ program for performing marker dependency filtering are also provided.

## Acknowledgements

The study was supported by American Heart Association Scientist Development Grant 13SDG17290032 (XY), Leducq Foundation (XY), American Heart Association Postdoctoral Fellowship 13POST17240095 (VPM), China Scholarship Council (LS), UCLA Eureka Scholarship (LS). MI and SB were supported by the Australian NHMRC (grant no. 1062227 & 1061435) and Australian Heart Foundation (grant no. 100038).

## Author contributions

XY and VPM conceived the study; VPM implemented the Mergeomics algorithms; LS performed the statistical analyses, method comparisons, and case studies; SB, TT, JK, SR, MI provided genetic data and conducted genetic association analysis; BZ provided support for KDA; LS, VPM and XY wrote the manuscript; SB, TT, JK, SR, MI reviewed and edited the manuscript.

